# Complement protein concentrations and activity in human cervical mucus

**DOI:** 10.64898/2026.07.18.739250

**Authors:** JG Marathe, E Mausser, JA Politch, M Tjilos, DJ Anderson

**Affiliations:** Department of Medicine, Boston University Aram V. Chobanian & Edward Avedisian School of Medicine, Boston, MA, USA; School of Public Health, Boston University, Boston, MA, USA

**Author notes:** Corresponding author: Jai G Marathe Department of Infectious Diseases, Boston Medical Center 670 Albany St. Suite 512 Boston, MA, USA 02118. These authors should be regarded as joint First Authors (JG Marathe, E Mausser).

**Keywords:** cervical mucus, complement, menstrual cycle, reproductive tract

## Abstract

**Problem:** Complement, a system of over 30 interacting proteins, functions as a critical immune mediator at mucosal surfaces including the intestine, airway, and nasal mucosae, where it orchestrates complement-dependent cytotoxicity (CDC), complement-dependent phagocytosis (CDP), and inflammatory responses. While complement components have been detected at low levels in genital tract fluids, including cervical mucus, the physiologic dynamics of complement in the female reproductive tract remain poorly characterized. Notably, systematic quantification of complement component levels across the menstrual cycle has not been conducted, limiting our understanding of how hormonal fluctuations may influence complement-mediated protection at the vaginal mucosa.

**Method of study:** Ten healthy women of reproductive age were recruited to provide paired samples of cervical mucus (CM) and serum (S) during each phase of the menstrual cycle: follicular, ovulatory, and luteal. Samples were analyzed using bead-based multiplex assays to quantify 13 primary complement components.

**Results:** All 13 complement proteins tested were detectable in CM and S; C3b/iC3b was the predominant complement component in CM followed by C4 and C3. In contrast, C4 was the dominant component in S followed by C1q and C3. There were significantly higher levels of C2 in CM during the follicular phase of the menstrual cycle vs. the ovulatory phase (1.15±0.80 vs 0.24±0.22µg/mL), C4b (0.56±0.35 vs 0.22±0.36 µg/mL), C5a (1.45±1.09 vs 0.38±0.48 µg/mL)

In contrast, no differences were found in serum complement levels were during the menstrual cycle. Complement concentrations were on average 308-fold (median= 84) lower in CM than in S, except for C3b/iC3b which was only 5 to 7.5-fold lower in CM, and C2 which was higher in CM at the luteal and follicular phases. Midcycle cervical mucus was capable of inducing complement-mediated hemolysis at about 1/3 the potency of serum.

**Conclusion:** Cervical mucus contains detectable and functional levels of complement proteins. These normative values can provide a foundation for future studies on immune mechanisms in the FRT.

## Introduction

The cervix is unique in its architecture and immunological function. It serves as a gateway to the uterus, allowing sperm passage during ovulation while preventing ascending infections through its mucus plug and tightly regulated opening^1^. The vagina hosts a unique microbiome dominated by Lactobacillus species, which produce lactic acid to maintain an acidic pH (around 3.5–4.5)^2,3^. This low pH creates an inhospitable environment for many pathogens, contributing to its protective function^4^. The cervix complements this protection by secreting mucus with antimicrobial properties, further inhibiting pathogen entry^1,5–7^.

Cervical mucus is an important immune mediator in the female reproductive tract (FRT) and has a role in both blocking movement of sperm as well as filtering and transporting sperm at ovulation^8–10^. It is primarily water but is also composed of lipids, cholesterols, carbohydrates, inorganic ions, glycans, and other proteins^1,10^.Cervical mucus undergoes cyclical changes in composition and consistency under the influence of hormones like estrogen and progesterone. In addition to containing immunoglobulins^11^, some complement proteins have been previously described in the cervical mucus^12–15^.

The complement system is composed of over thirty proteins that undergo a series of enzymatic steps to lyse and kill pathogenic cells^16^. These proteins are an integral part of the innate immune system of the cervix and the complement cascade is initiated by three distinct pathways: the classical, lectin, and alternative pathways^16^. All three pathways converge at the formation of C3 convertases. The classical pathway is triggered by immune complexes, primarily involving C1q binding to antibodies, triggering a cascade that leads to pathogen opsonization and lysis. The lectin pathway is similar but activated by mannose-binding lectin (MBL) recognizing carbohydrate structures on microbial surfaces, bypassing the need for antibodies^17^. The alternative pathway is continuously active at a low level and amplifies the immune response by direct recognition of pathogen surfaces, leading to the formation of the membrane attack complex (MAC) that lyses pathogens^18^. Together, these pathways provide a robust and versatile immune defense mechanism. In order for complement-mediated lysis to occur, all components of a cascade must be present as well as enzymatically functional.

Presence of complement C3 and its concentration in cervical mucus over the menstrual cycle was described as early as 1970^19^. Since then a few studies have evaluated the role of complement in infections like HIV^20^, pregnancy and parturition demonstrating that C4 levels are similar^21^ but C3 was lower during active labor compared to pregnancy^22^.

Complement regulation is important not only in the immune defense of the female reproductive tract but also in the protection of sperm and the trophoblast during conception and pregnancy^23^.The classical complement pathway is of great interest as it may be initiated by antibody-based drugs in the vaginal mucosa, both topically and systemically delivered. Manufactured antibodies can similarly interact with complement and contribute to Fc-mediated function, resulting in complement-mediated cell cytotoxicity. Anti-sperm antibodies have previously been shown to interact with serum complement and cause sperm immobilization and lysis, including Human Contraception Antibody (HCA)^24–26^ and a novel chimeric anti-gonococcal monoclonal antibody demonstrated increased efficacy due to complement activation in a mouse model^27^.

There is rapid growth in research evaluating Multipurpose Prevention Technologies (MPTs) designed to simultaneously prevent STIs, including HIV, and/or unintended pregnancy. Preclinical studies using a dissolvable film, MB66, containing two monoclonal antibodies (mAbs) against two sexually transmitted infections (STIs) showed HIV and HSV viral neutralization ex vivo 24 hours post-vaginal application^28^. The Human Contraception Antibody (HCA) is under investigation as a topical, nonhormonal, reversible contraceptive^24,28^. HCA, an antisperm antibody, may prevent pregnancy by rapidly and robustly agglutinating sperm when applied to the vaginal mucosa prior to intercourse^28^ and that sperm immobilization due to complement mediated cell-cytotoxicity may also contribute to its activity^29^. Complement proteins like C3, C4 remain the most commonly evaluated complement proteins in FRT but as additional complement proteins are described, there remains a gap in knowledge regarding their presence in cervical mucus over the course of the menstrual cycle. A broader investigation of complement concentrations is important for gaining insights into the immune landscape of the lower female reproductive tract, as well as to inform the development of antibody-based drugs for the female reproductive tract and other mucosal epithelia.

## Materials and methods

### Ethics statement

All studies were approved by the Institutional Review Board at Boston University Aram V. Chobanian & Edward Avedisian School of Medicine (Boston, MA, USA). Participants provided written informed consent prior to participation.

### Sample collection

#### Cervical mucus and serum

Ten healthy women of reproductive age 18-45 years and with a history of regular menstrual period were recruited to give paired cervical mucus and serum samples. Participants were excluded from the study if they were on hormonal birth control, unable to abstain from sexual intercourse for 48 hours prior to sample collection, had a prior history of genitourinary malignancy, pregnancy within 3 months of enrollment, douching within 48 hours of sample collection or were on chronic antibiotics. Three samples were collected from each participant: (1) 1-3 days after the end of their period (follicular phase), (2) within 48 hours after a positive ovulation test result (ovulatory phase), and (3) 3 days prior to the start of their next period (luteal phase). Samples were not required to be collected within a single menstrual cycle. An online ovulation calculator was used to estimate the date of ovulation for each participant^30^. The ovulation date for the participants was predicted by subtracting 14 days from the estimated last day of their menstrual cycle as per current clinical guidelines^31^. The participants were provided with an ovulation kit (Clearblue Digital Ovulation Predictor Kit, featuring Ovulation Test with digital results, Clearblue)^32^ and would use the kit starting 1-2 days before their calculated ovulation date. They would continue to test daily until they tested positive. The ovulatory samples were collected within 48 hours of a positive ovulation test.

At each visit, blood was collected by a trained technician into a red top vacutainer without additives (BD366408, BD, Franklin Lakes, NJ, USA). Next, an unlubricated speculum was inserted into the vagina and an endocervical pipelle (Aspirette Endocervical Pipelle, Cooper Surgical, Trumbull, CT, USA) was used to aspirate cervical mucus without contamination from vaginal secretions. Samples were processed immediately.

### Multiplex assays

Proteins in cervical mucus and serumwere quantified using commercial kits (complement: HCMP1MAG-19K and HCMP2MAG-19K, immunoglobulin isotypes: HGAMMAG-301K-06, hormones: MSHMAG-21K, Millipore Sigma, Burlington, MA, USA). From the ten female donors, 10 follicular, 8 ovulatory, and 10 luteal samples of serum and cervical mucus were assessed. All samples were kept on ice to prevent aberrant complement activation during the assay procedures. Plates were read on a Magpix System (Luminex Corporation, Austin, TX, USA). Purified human C3 (Cat# PHP283, reconstituted to 1 mg/mL Bio-Rad, Hercules, CA, USA,) was used to confirm sensitivity of assay and was used at dilutions of 1:10,000, 1:20,000, 1:40,000 and 1:80,000 to fit within the range of the standard curve. Serum was diluted following the manufacturer’s directions in the provided assay buffer. Cervical mucus dilutions varied by sample volume and predicted concentration. Follicular and luteal samples of cervical mucus were diluted to 1:10 or 1:1000 and ovulatory cervical mucus was diluted to either 1:5 or 1:500 for complement kit 1 or kit 2, respectively. For quantifying hormones, serum and cervical mucus, were extracted with acetonitrile, concentrated, and used neat. Some cervical mucus samples did not have sufficient volume for the assay, so any dilutions performed were accounted for during concentration calculations.

### Hemolysis

A standard hemolysis assay was performed^33,34^. Human red blood cells (RBCs) were isolated from whole blood by centrifugation and washed using triethanolamine-complement hemolysis buffer (TA-CHB). RBCs were spun at 900g for 5 minutes to pack the cells and were diluted to a 10% solution with TA-CHB. RBCs were sensitized using an equal volume of an anti-human RBC antibody at a final concentration of 3.4 mg/mL (Human Red Blood Cell RBC Antibody, Cat# 109-4139, RRID:AB_220010, Rockland Immunochemicals, Pottstown, PA, USA) by incubating in a water bath for 30 minutes at 30°C. The control complement source, human serum, underwent two-fold serial dilutions (final concentrations ranging from 1:2 to 1:32). Ovulatory cervical mucus was diluted 1:2 with 100μL of TA-CHB to improve miscibility (final dilution of 1:3). Heat inactivated (Hi) serum and cervical mucus (HiCM) were used as negative controls. Distilled water was used as a total lysis control and TA-CHB as a blank to account for any spontaneous lysis. For the assay, 100μL of sensitized RBCs were added to 100μL of each serum complement dilution, and 200μL was added for diluted cervical mucus samples. The solutions were incubated in a water bath at 37°C for 90 minutes with gentle agitation. The tubes were spun at 1500g for 5 minutes to pellet the RBCs, and 75μL of supernatant was added to 75μL of distilled water. The hemolytic activity was quantified by measuring the absorbance of the solution at 540nm on a plate reader. Percent lysis was calculated for both serum and cervical mucus by comparing the absorbance to the 100% lysis control. The experiment was run using midcycle cervical mucus from three different donors.

### Statistical analyses

The ovulation prediction data passed normality tests and was analyzed using a one sample t test with ∼14 days as the hypothetical value for ovulation day. Other data were log transformed and analyzed using a two-way ANOVA with a Tukey’s multiple comparison test. A test of sphericity (Geisser-Greenhouse’s epsilon) was used to determine the equality of variances of the pairwise differences of within subject conditions. Data was tested for normality prior to statistical analyses. Due to missing values, multiplex data was analyzed using a mixed-effects model with the Geisser-Greenhouse correction followed by Tukey’s multiple comparison test. Sperm immobilization data was analyzed using repeated measures two-way ANOVA also followed by Tukey’s multiple comparison test. GraphPad Prism (Version 10.0.0, GraphPad Software Inc., San Diego, CA, USA) was used for statistical analyses and graph creation. Differences were considered to be statistically significant when p <0.05.

## Results

### Cervical mucus and serum collection

The demographics of the 10 women recruited into the study can be seen in Table 1. The average age was 24.7±4.03 years. Five participants (50%) used condoms for birth control, two (20%) used a copper intrauterine device (IUD), and three (30%) did not use any birth control method. The average menstrual cycle frequency was 30.1±3.45 days with an average menses length of 6±0.82 days. Ovulatory samples could not be collected for two subjects: one never had a positive ovulation test over the course of the study and one discontinued participation. The actual day of ovulation varied from two days before to ten days after the estimated day of ovulation, an average of 2.07±3.10 days. The actual mean of ovulation day for the participants from onset of menses was -11.93 days and varied significantly (p=0.032) from the calculated mean (-14 days).

**Table 1.**
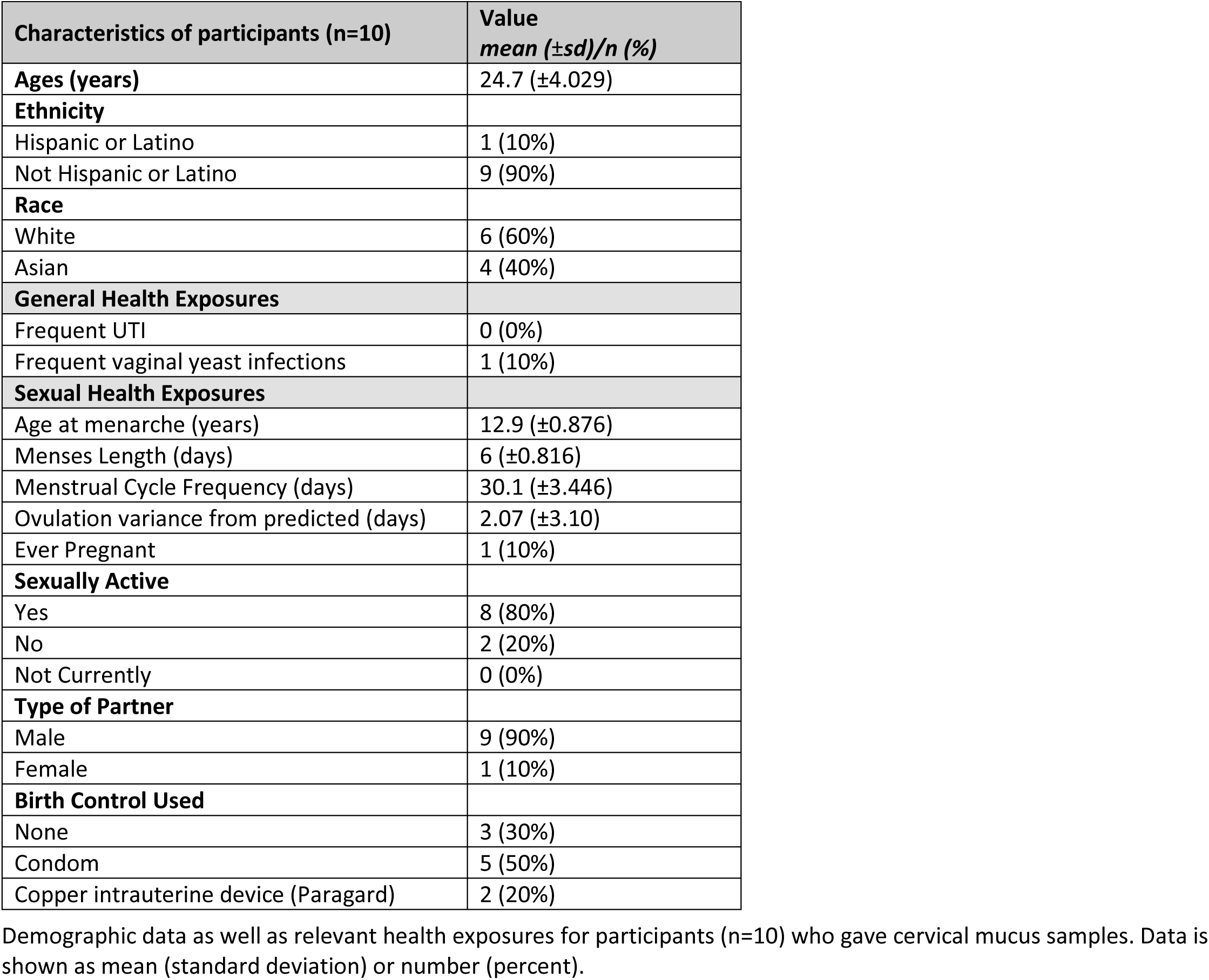
Demographics of female study participants.

### The effect of the menstrual cycle on complement levels in cervical mucus and serum

#### Changes in complement proteins across the menstrual cycle

The levels of 13 complement proteins were measured: C1q, C2, C3, C3b/iC3b, C4, C4b, C5, C5a, factor B, factor D (adipsin), factor H, factor I, and mannose-binding lectin (MBL). All of these complement components were detectable in every sample of cervical mucus and serum (Table 2). The menstrual cycle phase significantly influenced complement concentrations in cervical mucus, showing a substantial decline at ovulation compared to the follicular phase. Most of the complement components were present at reduced levels in cervical mucus at midcyle (ovulatory phase). Eight components exhibited a decrease of more than 50% in cervical mucus at midcycle compared to the follicular phase: C1q, C2, C4b, C5a, MBL, adipsin, factor H, and factor I; this difference was statistically significant for C2, C4b, C5a, and MBL (p < 0.05). Complement components often increased again during the luteal phase, (Figure 1).

**Figure 1:**
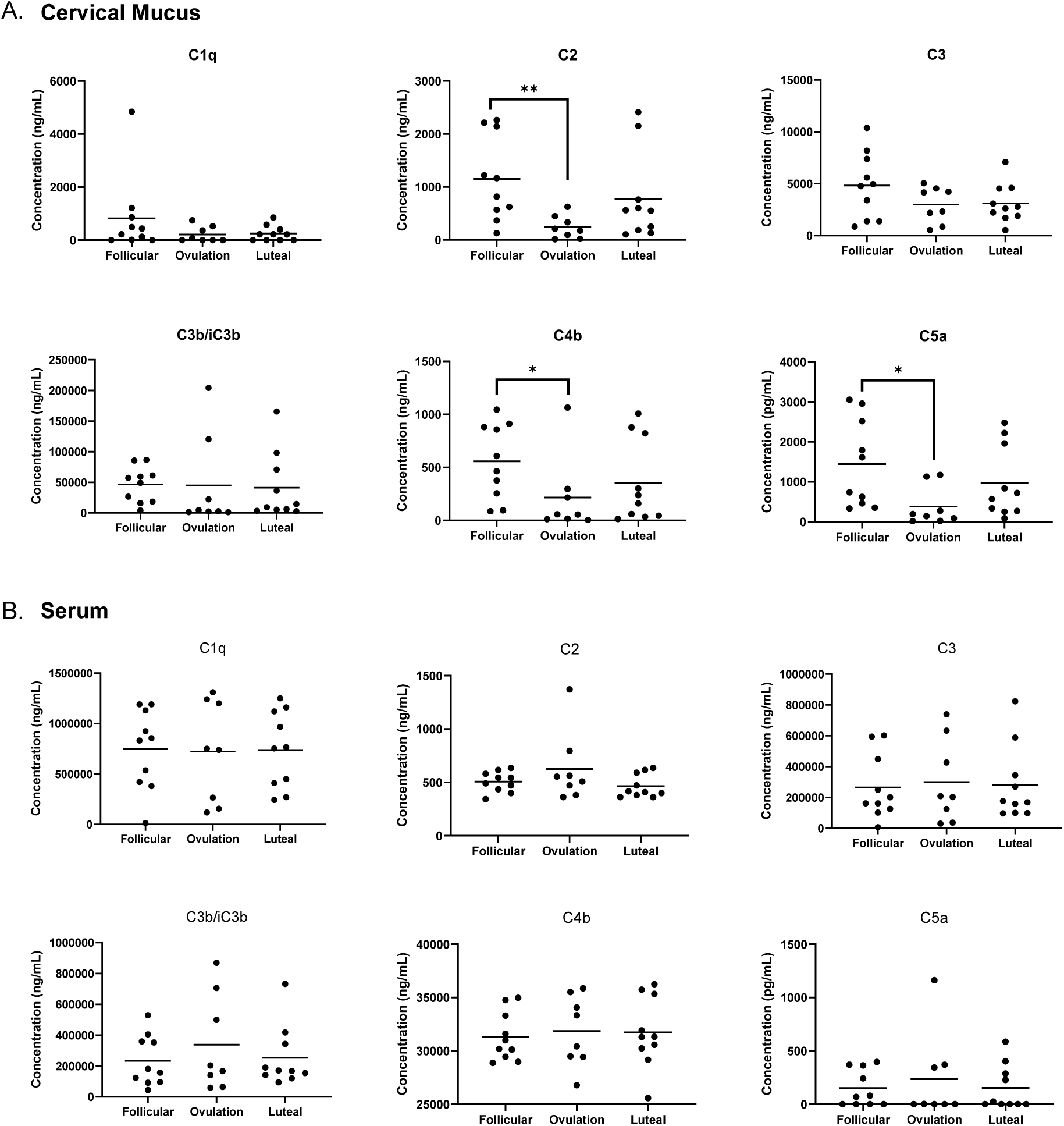
Concentrations of complement proteins of the classical pathway in cervical mucus (A) and serum (B) across menstrual cycle phases. Concentration of complement proteins C2, C4b and C5a was lower at ovulation in cervical mucus (*p<0.05, **p<0.005). There was no significant change in complement protein concentrations in serum across the menstrual phases.

**Table 2.** Differences in complement concentrations in cervical mucus and serum across the menstrual cycle.

|  |  | Cervical mucus |  |  | Serum |  |  |
| --- | --- | --- | --- | --- | --- | --- | --- |
|  | Menstrual Cycle Phase | Follicular (F) | Ovulation (O) | Luteal (L) | Follicular (F) | Ovulation (O) | Luteal (L) |
| Complement Pathway | Complement components | Value mean ±SD (μg/mL) | | | | | |
| Classical and Lectin Pathways | C1q | 0.82 ±1.47 | 0.21 ±0.30 | 0.25 ±0.29 | 746.89 ±396.47 | 722.15 ±498.34 | 738.60 ±379.52 |
| | MBL | 0.06 ±0.07 | 0.03 ±0.05** | 0.03 ±0.03 | 3.35 ±2.11 | 2.87 ±2.38 | 3.46 ±2.16 |
| | C2 | 1.15 ±0.80 | 0.24 ±0.22*** | 0.77 ±0.83 | 0.51 ±0.10 | 0.63 ±0.33 | 0.46 ±0.11 |
| | C4 | 22.04 ±21.38 | 12.88 ±17.65 | 14.58 ±19.71 | 3359.91 ±1352.09 | 3358.75 ±1587.45 | 3498.00 ±1012.85 |
| | C4b | 0.56 ±0.35 | 0.22 ±0.36* | 0.36 ±0.39 | 31.33 ±2.29 | 31.87 ±3.29 | 31.75 ±3.30 |
| Alternative Pathway | Factor B | 1.62 ±1.70 | 1.46 ±1.84 | 2.12 ±2.57 | 2076.03 ±1323.96 | 2387.49 ±1862.41 | 2245.24 ±1434.98 |
| | Factor D (Adipsin) | 0.12 ±0.15 | 0.04 ±0.03 | 0.05 ±0.04 | 2.89 ±0.77 | 2.58 ±0.57 | 2.43 ±0.62 |
| All Pathways | C3 | 4.83 ±3.18 | 2.98 ±1.74 | 3.10 ±1.87 | 264.77 ±209.42 | 299.89 ±270.03 | 281.95 ±243.02 |
| | C3b/iC3b | 46.39 ±28.96 | 44.85 ±76.15 | 41.20 ±54.47 | 233.22 ±164.27 | 338.18 ±312.30 | 252.73 ±196.58 |
| | C5 | 0.51 ±0.49 | 0.29 ±0.44 | 0.29 ±0.33 | 62.53 ±16.61 | 58.47 ±16.69 | 55.76 ±15.18 |
| | C5a | 0.002 ±0.001 | 0.0004 ±0.0005* | 0.001 ±0.001 | 0.0002 ±0.0002 | 0.00024 ±0.0004 | 0.0002 ±0.00021 |
| Inhibitors | Factor H | 13.12 ±11.20 | 4.83 ±4.50 | 6.78 ±6.61 | 2035.74 ±965.63 | 2065.27 ±106.38 | 2155.00 ±813.86 |
| | Factor I | 0.33 ±0.21 | 0.14 ±0.10 | 0.23 ±0.22 | 22.67 ±7.93 | 19.74 ±6.11 | 20.544 ±6.64 |

Cervical mucus and serum samples are stratified by menstrual cycle phase: follicular, ovulation, and luteal. C3b/iC3b and C4 represent the largest fraction of complement proteins across all menstrual phases for cervical mucus.

In serum, complement concentrations did not show any significant fluctuations over the course of the menstrual cycle (Table 2). The large standard deviations for complement concentrations were due to substantial interpatient variability, but intrapatient variability across all three visits was low across all three visits. Complement levels in cervical mucus were on average 308-fold (median = 84.02) lower than serum but the fold ratio of serum to cervical mucus varied from 0.2 for C5a to 1716.95 for C1q.

As seen in (Figure 2), the distribution of classical complement components also varied between serum and cervical mucus. In serum, the most abundant protein was C4, followed by C1q, C3 and C3b/iC3b. These distributions stayed relatively consistent in the follicular, ovulatory, and luteal samples. Conversely, the most abundant complement proteins in cervical mucus were C3b/iC3b, C4, and C3. The total concentration of classical complement components decreased from the follicular to the ovulatory samples and stayed low in the luteal phase. As seen previously, this was a result of a decrease in most of the components, most clearly seen with C4 decreasing from 22 μg/mL to 12 μg/mL.

**Figure 2:**
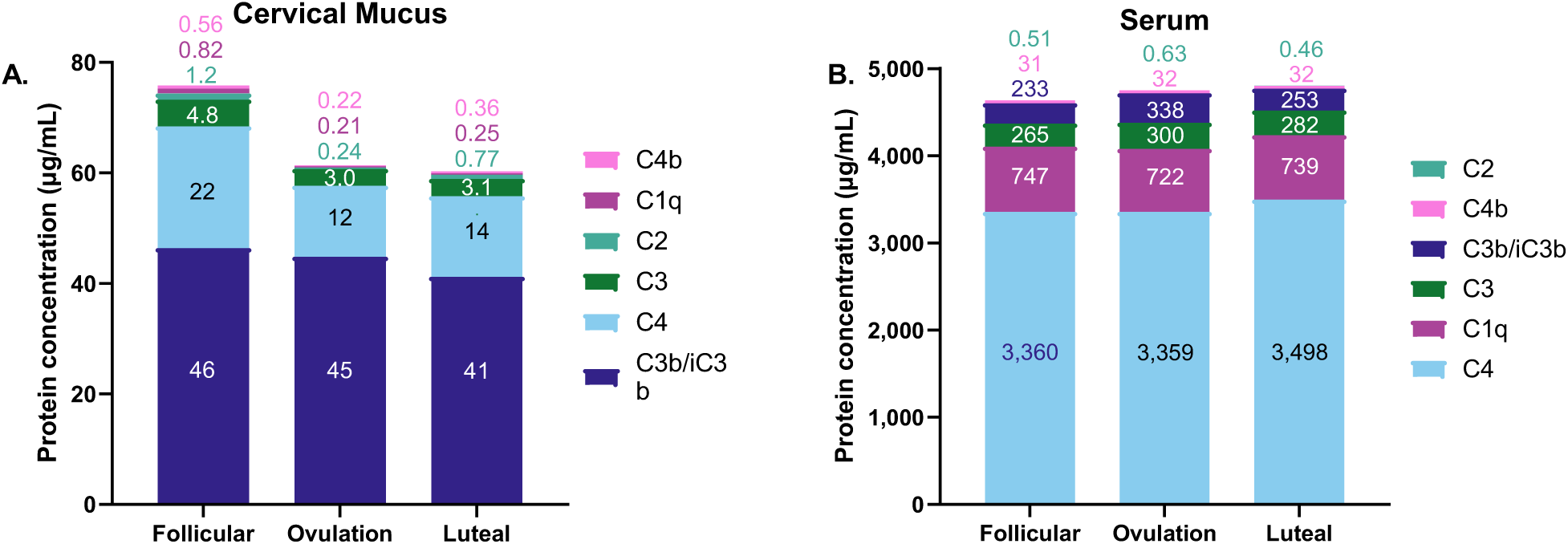
Concentrations of the major classical complement proteins in cervical mucus (A) and serum (B) at different phases of the menstrual cycle. Each bar is labeled with the concentration of that component within or above the bar. C3b/iC3b predominated in cervical mucus whereas C4 was the most abundant complement protein in serum across all phases of the menstrual cycle.

#### Hormone levels

The levels of hormones in cervical mucus were lower than those detected in serum and did not show statistically significant differences across the menstrual cycle (Supplement table 1). The levels of hormones in serum were within the range of normal serum laboratory values from previous studies^35^. In serum, estradiol was low during the follicular phase (0.085±0.05 ng/mL) and increased to peak at ovulation (0.22±0.16 ng/mL, p=0.048) corroborating the positive ovulation tests. Progesterone was low during the follicular phase (5.44±2.76 ng/mL), and ovulatory phase (8.91±4.48 ng/mL), and peaked at the luteal phase to 13.04±9.48 ng/mL (p <0.05).

### Comparison of Complement-mediated hemolytic activity of cervical mucus and serum

As cervical mucus was found to contain all 13 of the evaluated proteins in the complement cascade, the functionality of complement was tested with the hemolysis assay in 3 donors.

The hemolytic activity of cervical mucus varied amongst the donors, and was 7.4%, 8.4% and 49.4%; on average, cervical mucus had 10.6% the hemolytic activity of serum. (Figure 3).

**Figure 3:**
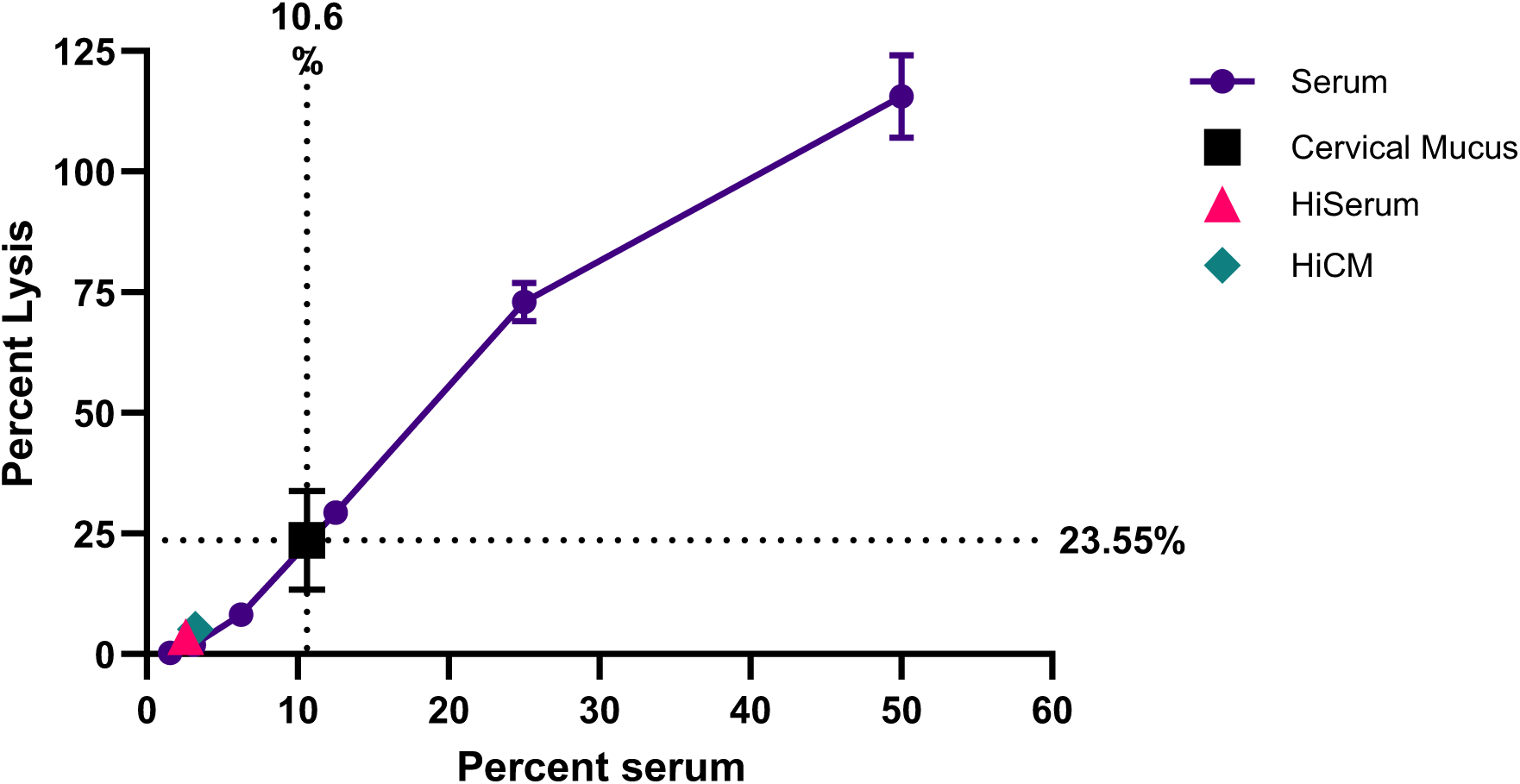
Cervical mucus lysed 23.55% of RBCs after treatment with anti-RBC antibodies, this is the equivalent of the hemolytic activity of serum at a ∼10% dilution. Heat inactivated (Hi) Serum and CM were used as negative controls and showed minimal RBC lysis.

## Discussion

The complement system plays a key immune defense role at mucosal surfaces. In reproductive mucosa, hormonal variation modulates the immune functions^36^. Here, we determine that complement levels decrease during ovulation. This reduction likely helps prevent strong immune responses to sperm, thereby increasing the chances of successful conception. We found that concentrations of all thirteen components of the complement cascade measured in this study decreased between the follicular and ovulatory phases, supporting this theory. The local production of complement has been shown to be hormonally regulated, stimulated by estrogen and inhibited by progesterone^37,38^, which could explain the mechanism for the temporal variations in cervical mucus observed in this study. We also report that despite the lower concentrations, complement in ovulatory cervical mucus was functional and able to lyse target cells similar to and earlier finding by Price and Boettcher^39^. Differences in serum complement concentrations over the phases of menstrual cycle were not seen, indicating that these changes are restricted to the FRT.

The immune microenvironment of the female tract changes around ovulation possibly to be more receptive to sperm by increasing permeability of cervical mucus and decreasing the levels of immune components^19,35,40^. In this “trade-off paradigm,” the immune responses in the FRT must be robust enough to prevent infection but must also be regulated to allow for fertilization and fetal development. A decrease in complement levels at ovulation would facilitate this function. The varied composition of complement components between serum and cervical mucus, suggests different functions for each of these biological fluids. Potential functions of complement in cervical mucus include complement-dependent cytotoxicity (CDC) of pathogens and sperm either in the presence of antibody^29,41^ or membrane damage^42^ , opsonization and complement-dependent phagocytosis (CDP), and induction of inflammation (refs).

Complement components are largely produced in the liver^23^, although there is evidence that C3 is also locally produced In the endometrium^43^. Differences in complement levels in cervical mucus at midcycle is likely due to dilution effects of secreted mucins. The complosome, referring to the intracellular complement system, is being increasingly recognized as playing a role locally within tissues by regulating critical cellular processes including metabolism, gene transcription, autophagy^44^. Complosome activation was described in vaginal cells; it was shown to have differential activation during vulvovaginal candidiasis indicating a role in activation and resolution of inflammation^45^.

In the upper FRT, complement is hypothesized to play a role in fertilization. The outer membrane of sperm is coated in anti-complement factors like CD59 and decay accelerating factor (DAF)^46^ . Membrane cofactor protein (MCP/CD46) is present on the inner acrosomal membrane, so it is only exposed following capacitation. This has led to the theory that MCP is involved in sperm-oocyte fusion. This hypoglycosylated sperm-specific variant of MCP can bind dimeric C3b which could link the sperm to the oocyte via binding to complement receptor type 1 (CD35) or type 3 (CD11b/CD18)^47,48^. Complement in the FRT is typically studied in the context of pregnancy and has also been reported to play a role in labor^49^ . C3 has been shown to be immunoprotective to the trophoblast and C1q may be an embryotrophic factor^50,51^. Complement may also have additional novel functions in the lower FRT and at other sites^52–54^. Livson et. al,. reported high levels of C3 in cervical secretions and C3 activation in non-pregnant and pregnant women and also during labor^22^. C3 activation products were significantly higher during pregnancy and decreased during labor indicating role for C3 activation during pregnancy. Our data demonstrate that there are high levels of C3b/iC3b throughout the menstrual cycle and that these levels do not fluctuate significantly. The presence of C3b/iC3b as the largest fraction of complement in lower FRT may suggest the dynamic interplay between host innate immune response and the local microbiome. This fraction, which is activated through all three complement pathways, acts as an opsonin and we hypothesize that it may be important in maintaining the homeostasis in the lower FRT.

Most studies characterizing the proteome of cervical mucus have used cervicovaginal lavages (CVLs) in which the cervix and vagina were washed with saline. The resulting samples can be concentrated, but it is extremely difficult to determine the original concentration of proteins due to variable volumes. Cervical or vaginal secretions collected with swabs or gauze can be weighted prior to elution of the sample from the collection matrix and therefore provide more accurate measurements. By collecting cervical mucus directly, we were able to minimize contamination with vaginal secretions and cells and obtain measurable volumes. However, due to its strong viscoelastic properties, the cervical mucus samples required dilution in order to pipette sample volumes accurately. Therefore, we were unable to use neat cervical mucus in experiments but obtained actual concentrations by correcting for the dilution. As cervical secretions increase around ovulation, larger volumes of midcycle mucus were collected. It has been suggested that the absolute amount of proteins in cervical mucus remains constant but decreased concentrations are observed at ovulation due to the increased volume due to the secretion of mucus^55,56^. There was insufficient volume of follicular and luteal samples to test in the hemolytic functional test, but we hypothesize that these samples would have comparable or greater hemolytic function due to the higher concentrations of complement.

As serum complement (particularly C3 and C4) are well established laboratory values, we were surprised that the serum concentrations measured in this study deviated from clinical laboratory values; however, this is consistent with findings reported by other research studies^57,58^ and is likely be an assay-based difference.

When developing antibodies for use in the female reproductive tract (FRT), complement-dependent cytotoxicity (CDC) targeting sperm or pathogens could provide additional protection against unintended pregnancy and sexually transmitted infections. For the first time, the majority of complement components have been identified in human cervical mucus, where the classical complement cascade appears functional. This system can lyse red blood cells in the presence of RBC antibodies, albeit less efficiently than serum. As part of the innate immune system, complement may contribute to the clearance of sexually transmitted pathogens in the FRT.

Currently, our group is developing a human monoclonal antibody, the human contraceptive antibody (HCA), as a contraceptive^24^. While its primary mechanism involves Fab-mediated sperm agglutination, HCA may also have additional Fc-mediated functions in the FRT. Previous studies have demonstrated that HCA, through Fc interactions, can immobilize and lyse sperm in the presence of a complement source and trap sperm in ovulatory cervical mucus^26^. The interaction between monoclonal antibodies and complement in cervical mucus may enhance antibody efficacy; we recently reported results from a postcoital test involving the application of HCA film prior to intercourse that complement-mediated cytotoxicity was likely a major mechanism immobilizing sperm in cervical mucus. Anderson D Fert Steril). However, it remains unclear whether these interactions at mucosal surfaces also trigger proinflammatory responses which could promote the transmision of HIV and potentially other pathogens.; this would be important to evaluate during future clinical trials. Overall, the complement proteins in the reproductive tract are an understudied but important component of the mucosal immune system. This study provides normative values that might promote the development of novel monoclonal antibodies targeting sperm and pathogens the female reproductive tract.

## Data availability statement

The data underlying this article are available in the article and in its online supplementary material.

## Acknowledgements

We would like to thank our study participants and the staff at the General Clinical Research Unit (GCRU) at Boston University for use of their facilities and assistance. Our thanks to J Matt Au for assisting with the multiplex instrumentation. This work was supported by the Boston University Analytical Instrumentation Core.

## Authors’ roles

JGM conceived of the study, collected patient samples, and assisted with sample preparation. EM performed quantification and functional assays. JAP assisted in data analysis and statistical calculations. MT helped with participant recruitment and scheduling visits. All authors assisted with manuscript preparation.

## References

1. Han L, Taub R, Jensen JT. Cervical mucus and contraception: what we know and what we don’t. Contraception. 2017;96(5):310–21.

2. Boskey ER, Cone RA, Whaley KJ, Moench TR. Origins of vaginal acidity: high D/L lactate ratio is consistent with bacteria being the primary source. Hum Reprod. 2001;16(9):1809–13.

3. Boskey ER, Telsch KM, Whaley KJ, Moench TR, Cone RA. Acid production by vaginal flora in vitro is consistent with the rate and extent of vaginal acidification. Infect Immun. 1999;67(10):5170–5.

4. O’Hanlon DE, Moench TR, Cone RA. Vaginal pH and microbicidal lactic acid when lactobacilli dominate the microbiota. PLoS One. 2013;8(11):e80074.

5. Hein M, Valore EV, Helmig RB, Uldbjerg N, Ganz T. Antimicrobial factors in the cervical mucus plug. Am J Obstet Gynecol. 2002;187(1):137–44.

6. Ming L, Xiaoling P, Yan L, Lili W, Qi W, Xiyong Y, et al. Purification of antimicrobial factors from human cervical mucus. Human reproduction. 2007;22(7):1810–5.

7. Domino SE, Hurd EA, Thomsson KA, Karnak DM, Holmén Larsson JM, Thomsson E, et al. Cervical mucins carry α (1, 2) fucosylated glycans that partly protect from experimental vaginal candidiasis. Glycoconjugate journal. 2009;26:1125–34.

8. Katz DF, Slade DA, Nakajima ST. Analysis of pre-ovulatory changes in cervical mucus hydration and sperm penetrability. Adv Contracept. 1997;13(2-3):143–51.

9. Yudin AI, Hanson FW, Katz DF. Human cervical mucus and its interaction with sperm: a fine-structural view. Biol Reprod. 1989;40(3):661–71.

10. Lacroix G, Gouyer V, Gottrand F, Desseyn JL. The Cervicovaginal Mucus Barrier. Int J Mol Sci. 2020;21(21).

11. Schumacher GF. Immunology of spermatozoa and cervical mucus. Hum Reprod. 1988;3(3):289–300.

12. Clark GF, Schust DJ. Manifestations of immune tolerance in the human female reproductive tract. Front Immunol. 2013;4:26.

13. Cone RA. Barrier properties of mucus. Adv Drug Deliv Rev. 2009;61(2):75–85.

14. Schumacher G. Hormonal immune factors in the female reproductive tract and their changes during the cycle. Immunological aspects of infertility and fertility regulation. 1980:93–141.

15. Shaw JL, Smith CR, Diamandis EP. Proteomic analysis of human cervico-vaginal fluid. Journal of proteome research. 2007;6(7):2859–65.

16. Sarma JV, Ward PA. The complement system. Cell Tissue Res. 2011;343(1):227–35.

17. Wallis R. Interactions between mannose-binding lectin and MASPs during complement activation by the lectin pathway. Immunobiology. 2007;212(4-5):289–99.

18. Qu H, Ricklin D, Lambris JD. Recent developments in low molecular weight complement inhibitors. Mol Immunol. 2009;47(2-3):185–95.

19. Schumacher GF. Biochemistry of cervical mucus. Fertil Steril. 1970;21(10):697–705.

20. Beverly ES, D’Amico RD, Landay AL, Spear GT, Massad LS, Rydman RJ, et al. Evaluation of immunologic markers in cervicovaginal fluid of HIV-infected and uninfected women: implications for the immunologic response to HIV in the female genital tract. JAIDS Journal of Acquired Immune Deficiency Syndromes. 1997;16(3):161–8.

21. Livson S, Virtanen S, Lokki AI, Holster T, Rahkonen L, Kalliala I, et al. Cervicovaginal complement activation and microbiota during pregnancy and in parturition. Frontiers in immunology. 2022;13:925630.

22. Livson S, Jarva H, Kalliala I, Lokki AI, Heikkinen-Eloranta J, Nieminen P, et al. Activation of the Complement System in the Lower Genital Tract During Pregnancy and Delivery. Front Immunol. 2020;11:563073.

23. Vanderpuye OA, Labarrere CA, McIntyre JA. The complement system in human reproduction. Am J Reprod Immunol. 1992;27(3-4):145–55.

24. Baldeon-Vaca G, Marathe JG, Politch JA, Mausser E, Pudney J, Doud J, et al. Production and characterization of a human antisperm monoclonal antibody against CD52g for topical contraception in women. EBioMedicine. 2021;69:103478.

25. D’Cruz OJ, Haas GG, Jr., Wang BL, DeBault LE. Activation of human complement by IgG antisperm antibody and the demonstration of C3 and C5b-9-mediated immune injury to human sperm. J Immunol. 1991;146(2):611–20.

26. Mausser E, Nador E, Politch JA, Pauly MR, Marathe JG, Moench TR, et al. LALAPG variant of the Human Contraception Antibody (HCA) reduces Fc-mediated effector functions while maintaining sperm agglutination activity. PLoS One. 2023;18(3):e0282147.

27. Gulati S, Beurskens FJ, de Kreuk BJ, Roza M, Zheng B, DeOliveira RB, et al. Complement alone drives efficacy of a chimeric antigonococcal monoclonal antibody. PLoS Biol. 2019;17(6):e3000323.

28. Thurman AR, Moench TR, Hoke M, Politch JA, Cabral H, Mausser E, et al. ZB-06, a vaginal film containing an engineered human contraceptive antibody (HC4-N), demonstrates safety and efficacy in a phase 1 postcoital test and safety study. Am J Obstet Gynecol. 2023;228(6):716 e1-e12.

29. Nador E, Mausser E, Marathe JG, Politch JA, Thurman AR, Whaley KJ, et al. Antibody-dependent complement-mediated sperm cytotoxicity in the endocervix is a dominant contraceptive mechanism of ZB-06 vaginal film. Fertil Steril. 2024.

30. Office on Women’s Health in the U.S. Department of Health and Human Services. Ovulation calculator [cited 2025 2/7/2025]. Available from: https://www.womenshealth.gov/ovulation-calculator.

31. Wilcox AJ, Dunson D, Baird DD. The timing of the "fertile window" in the menstrual cycle: day specific estimates from a prospective study. BMJ. 2000;321(7271):1259–62.

32. Behre HM, Kuhlage J, Gassner C, Sonntag B, Schem C, Schneider HP, et al. Prediction of ovulation by urinary hormone measurements with the home use ClearPlan Fertility Monitor: comparison with transvaginal ultrasound scans and serum hormone measurements. Hum Reprod. 2000;15(12):2478–82.

33. Inglis JE, Radziwon KA, Maniero GD. The serum complement system: a simplified laboratory exercise to measure the activity of an important component of the immune system. Advances in Physiology Education. 2008;32(4):317–21.

34. Costabile M. Measuring the 50% haemolytic complement (CH50) activity of serum. Journal of visualized experiments: JoVE. 2010(37):1923.

35. Schumacher GF, Kim MH, Hosseinian AH, Dupon C. Immunoglobulins, proteinase inhibitors, albumin, and lysozyme in human cervical mucus. I. Communication: hormonal profiles and cervical mucus changes--methods and results. Am J Obstet Gynecol. 1977;129(6):629–36.

36. Nguyen PV, Kafka JK, Ferreira VH, Roth K, Kaushic C. Innate and adaptive immune responses in male and female reproductive tracts in homeostasis and following HIV infection. Cellular & molecular immunology. 2014;11(5):410–27.

37. Li SH, Huang HL, Chen YH. Ovarian steroid-regulated synthesis and secretion of complement C3 and factor B in mouse endometrium during the natural estrous cycle and pregnancy period. Biol Reprod. 2002;66(2):322–32.

38. Lundeen SG, Zhang Z, Zhu Y, Carver JM, Winneker RC. Rat uterine complement C3 expression as a model for progesterone receptor modulators: characterization of the new progestin trimegestone. J Steroid Biochem Mol Biol. 2001;78(2):137–43.

39. Price RJ, Boettcher B. The presence of complement in human cervical mucus and its possible relevance to infertility in women with complement-dependent sperm-immobilizing antibodies. Fertil Steril. 1979;32(1):61–6.

40. Fleetwood L, Landgren BM, Eneroth P. A method for isolation, characterization, and quantitation of soluble proteins in uterine cervical secretion. Gynecol Obstet Invest. 1984;17(1):40–6.

41. Vogelpoel FR, te Velde ER, Scheenjes E, Van Kooy R, Kremer J, Verhoef J. Antibody and complement-binding activity of viable and nonviable human spermatozoa. Arch Androl. 1987;18(3):189–97.

42. Dohadwala S, Shah P, Farrell MK, Politch JA, Marathe J, Costello CE, et al. Sialidases derived from Gardnerella vaginalis and Prevotella timonensis remodel the sperm glycocalyx and impair sperm function. Glycobiology. 2025;35(11).

43. Bischof P, Planas-Basset D, Meisser A, Campana A. Immunology: Investigations on the cell type responsible for the endometrial secretion of complement component 3 (C3). Human Reproduction. 1994;9(9):1652–9.

44. Freiwald T, Afzali B. The secret life of complement: challenges and opportunities in exploring functions of the complosome in disease. J Clin Invest. 2025;135(12).

45. Kenno S, Pedretti N, Spaggiari L, Ardizzoni A, Comar M, Posch W, et al. Vaginal Clinical Isolates of Candida albicans Differentially Modulate Complosome Activation in Vaginal Epithelial Cells. J Fungi (Basel). 2025;11(7).

46. Morgan B, Harris C. Complement regulation in the reproductive system. Complement regulatory proteins San Diego: Academic Press. 1999:192-206.

47. Anderson DJ, Abbott AF, Jack RM. The role of complement component C3b and its receptors in sperm-oocyte interaction. Proc Natl Acad Sci U S A. 1993;90(21):10051–5.

48. Riley-Vargas RC, Lanzendorf S, Atkinson JP. Targeted and restricted complement activation on acrosome-reacted spermatozoa. J Clin Invest. 2005;115(5):1241–9.

49. Girardi G, Lingo JJ, Fleming SD, Regal JF. Essential Role of Complement in Pregnancy: From Implantation to Parturition and Beyond. Front Immunol. 2020;11:1681.

50. Kouser L, Madhukaran SP, Shastri A, Saraon A, Ferluga J, Al-Mozaini M, et al. Emerging and Novel Functions of Complement Protein C1q. Front Immunol. 2015;6:317.

51. Usami M, Mitsunaga K, Miyajima A, Sunouchi M, Doi O. Complement component C3 functions as an embryotrophic factor in early postimplantation rat embryos. Int J Dev Biol. 2010;54(8-9):1277–85.

52. Hawksworth OA, Coulthard LG, Mantovani S, Woodruff TM. Complement in stem cells and development. Semin Immunol. 2018;37:74–84.

53. King BC, Kulak K, Krus U, Rosberg R, Golec E, Wozniak K, et al. Complement Component C3 Is Highly Expressed in Human Pancreatic Islets and Prevents beta Cell Death via ATG16L1 Interaction and Autophagy Regulation. Cell Metab. 2019;29(1):202–10 e6.

54. Giclas PC, King TE, Baker SL, Russo J, Henson PM. Complement activity in normal rabbit bronchoalveolar fluid. Description of an inhibitor of C3 activation. Am Rev Respir Dis. 1987;135(2):403–11.

55. Moghissi K, FN S. Cyclic changes in the amount and sialic acid of cervical mucus. 1976.

56. Ozdian T, Vodicka J, Dostal J, Holub D, Vaclavkova J, Jeseta M, et al. Proteome Mapping of Cervical Mucus and Its Potential as a Source of Biomarkers in Female Tract Disorders. Int J Mol Sci. 2023;24(2).

57. Keith R, Gilliam B, Pepin D, Xiao Q. Complement factor serum and plasma levels measured by a MILLIPLEX((R)) Magnetic Bead Panel: A reply. Scand J Immunol. 2021;93(6):e13036.

58. Sandholm K, Carlsson H, Persson B, Skattum L, Tjernberg I, Nilsson B, et al. Discrepancies in plasma levels of complement components measured by a newly introduced commercially available magnetic bead technique compared to presently available clinical reference intervals. Scand J Immunol. 2020;91(2):e12831.

