## Supplemental Table 1 for "Complement protein concentrations and activity in human cervical mucus"

Supplement Table 1: **Differences in hormone concentrations in serum, and cervical mucus across the menstrual cycle.**

| | **Menstrual Cycle Phase** | **Follicular (F)** | **Ovulation (O)** | **Luteal (L)** | **Follicular (F)** | **Ovulation (O)** | **Luteal (L)** |
|---|---|---|---|---|---|---|---|
| | | **Cervical Mucus** | | | **Serum** | | |
| **Hormones (ng/mL)** | Cortisol | 5.83 ±5.58 | 3.66 ±2.11 | 5.30 ±5.94 | 62.08 ±8.77 | 70.05 ±14.42 | 68.14 ±13.16 |
| | Estradiol | 0.03 ±0.02 | 0.04 ±0.03 | 0.04 ±0.03 | 0.085 ±0.05 | 0.22 ±0.16 | 0.16 ±0.12 |
| | Progesterone | 0.18 ±0.04 | 0.19 ±0.00 | 0.24 ±0.11 | 5.44 ±2.76*** | 8.91 ±4.48**** | 13.04 ±9.48****** |
| | Testosterone | 0.15 ±0.9 | 0.15 ±0.12 | 0.14 ±0.08 | 0.77 ±0.32 | 0.79 ±0.31 | 0.73 ±0.30 |
| | T3 | 0.42 ±0.16 | 0.31 ±0.13 | 0.38 ±0.15 | 2.17 ±0.89 | 2.05 ±0.27 | 2.05 ±0.40 |
| | T4 | 1.74 ±1.05 | 1.10 ±0.87 | 1.36 ±1.45 | 60.69 ±13.03 | 55.80 ±9.13 | 61.75 ±8.75 |

Serum and cervical mucus samples are stratified by menstrual cycle phase: follicular, ovulation, and luteal. Hormones are in ng/mL. (*$p < 0.05$ F vs O, **$p < 0.05$ F vs L and ***$p < 0.05$ L vs O).
